# Synthetic adjuvants to potentiate the immune response against rAg85B by nasal route

**DOI:** 10.64898/2026.08.31.748287

**Authors:** Rong Chen, Changyuan Tan, Wen Li, Yasser Perera, Yadira Lobaina

## Abstract

Respiratory infections remain a relevant global health concern nowadays. The rapid dissemination and high mutational ratio of some pathogens, amid human demographic drivers, increases their pandemic potential. On the other hand, some bacterial infections have re-emerged with new resistant variants, supporting the need of updated or more potent vaccines. The development of nasal vaccines, with the capacity to efficiently induce immune response at the respiratory tract and systemic compartments, constitutes an appealing strategy to tackle respiratory infections, including tuberculosis. In this work, two synthetic compounds, the ODN-39M and the peptide LALF_32-51_, are evaluated as nasal vaccine adjuvants for the Ag85B antigen from Mycobacterium tuberculosis. The capacity of such adjuvants to enhance the immune response at systemic and mucosal compartments following intranasal immunization was assessed in Balb/C mice. The ODN-39M was able to potentiate the immune response, showing preferentially a Th1 pattern. The vaccine preparation containing this CpG adjuvant plus Ag85B induces potent cell-mediated and high IgA specific antibody response in lungs, along with systemic immunity. A preliminary study of the interaction between the ODN-39M with the NALT immune cells shows the early recruitment of lymphoid cells, and the increase of CD69 activation marker, mainly in B and dendritic cells. Although further studies should be done to reach a deeper understanding of NALT functioning, including the development of local innate response, the current data enrich the scarce reports focused on this compartment and support the use of the ODN-39M as a potent and potentially safe adjuvant option for future nasal vaccines.

## 1. Introduction

Among relevant respiratory infections, Tuberculosis (TB), caused by Mycobacterium tuberculosis (Mtb), remains as a major health problem affecting over one-third population of the world. Estimates, previous to COVID-19 pandemic, indicate that TB is one of the world’s deadliest infectious diseases, with more than 1.3 million deaths per year, mostly in low- and middle-income countries [**1**].

So far, Bacillus Calmette–Guerin (BCG), a vaccine dating from almost 100 years ago, constitutes the only licensed vaccine against TB. BCG is administered after birth and can edectively protect infants from severe forms of infection, but fails in protect adolescents and young adults from pulmonary TB [**2, 3**]. This fact contributes to sustain a pool of Mtb carriers that account for infection transmission [**3**]. In general, BCG-induced protection is variable and the generated immunity last between 10-20 years. Re-vaccination with this vaccine is not recommended by WHO, considering that booster doses do not provide much additional protection. Furthermore, among the more susceptible populations to develop pulmonary TB are the immunocompromised individuals, especially the HIV-infected. For this growing group, including HIV positive infants, the administration of BCG is not recommended considering the nature of the vaccine preparation based on live attenuated bacteria [**4**]. In addition, in the past years the rise of drug-resistant and multiple drug-resistant strains of Mtb adds further complexity to the landscape of TB prophylaxis and treatment [**5**].

Altogether, the current situation reinforces the need of new vaccines against TB. Nowadays several candidates are under evaluation at preclinical and early clinical stages. The majority of the vaccine candidates in study use parenteral administration routes and are based on attenuated bacteria or whole cell extracts. Those few that are nasally delivered are based on adenoviral vector platform [**6–8**]. Conversely, our team strongly support the development of protein subunit-based nasal vaccine candidates against respiratory pathogens. In this line we have recently published some works related with the development of broad-spectrum anti-coronavirus vaccine candidates [**9–11**].

The use of mucosal delivery for vaccines administration was traditionally neglected, with only few successfully approved vaccines [**12**]. However, after COVID-19 pandemic and supported by the current advances in related sciences (immunology, genomics, vaccinology) this strategy has been reinvigorated. Despite the advantages, it is recognized that mucosal administration imposes some challenges to the drug/vaccine formulations, related with the physiological role and features of these surfaces [**12, 13**]. In the specific case of vaccine preparations, the use of delivery systems, nanoparticulate structures, and adjuvants is crucial to generate a proper immune response [**14, 15**]. To date, the more potent known mucosal adjuvants are natural toxins (like Vibrio cholerae and *E. coli* toxins) and derivatives, which imply an obvious safety concern, limiting their use in human [**16**]. Consequently, the search for novel potent, safe, and cost-effective adjuvants for intranasal delivery of vaccines remains as a relevant research subject nowadays.

Among the arsenal of new mucosal adjuvants, the CpG ODNs has shown potential as potent activators of antigen presenting cells, through TLR9 stimulation, and strong inducers of Th1-biased immune response. Specifically, the ODN-39M has been evaluated previously, as component of different vaccine candidates, by parenteral and nasal route with promising results in mice and monkeys [**9–11, 17–20**]. On the other hand, the cell-penetrating peptides (CPPs) has also been tested as immune-enhancers by nasal route with appealing results [**21–23**]. In the current work we explored the nasal immunogenicity of vaccine preparations based on Ag85B protein, one of the most relevant TB target antigens, and two different synthetic adjuvants. In addition, preliminary studies on interaction between adjuvants and the nasal-associated lymphoid tissue (NALT) immune cells were also carried out to elucidate the mechanisms of action associated with their local effect at nasal mucosa.

## 2. Materials and Methods

### 2.1. Biological reagents

The recombinant Mycobacterium tuberculosis Diacylglycerol acyltransferase/mycolyltransferase Ag85B (fbpB) antigen was purchased from Cusabio Technology LLC (China). The protein (CSB-EP314366MVZa0), containing a N-terminal 6xHis-tag label, was produced in *E. coli* with 94% purity.

The ODN-39M, a 39 mer, whole phosphodiester backbone oligodeoxynucleotide (5’-ATC GAC TCT CGA GCG TTC TCG GGG GAC GAT CGT CGG GGG-3’), was synthesized by Sangon Biotech (China).

The peptide LALF_32-51_ (HYRIKPTFRRLKWKYKGKFW), C-terminal amidated, was synthesized with > 95% purity by Sangon Biotech (China).

### 2.2. Immunization experiments

Beijing Vital River Laboratory Animal Technology Co., Ltd. conducted the mice experiments. The animal facility complies with the national standard of the People’s Republic of China GB14925-2010. The immunization protocols were approved by Institutional Animal Care and Use Committee. Six to eight weeks old, females, Balb/C (inbred, H-2d) mice were employed.

Balb/c mice were distributed in four groups of five animals each and immunized with three doses, administered on days 0, 15 and 30. Four treatment groups were intranasally immunized: G1(10µg Ag85B), G2 (10µg Ag85B + 2mM LAL_32-51_), G3 (10µg Ag85B + 20µg ODN-39M), and G4 (PBS 1X). All immunogens were dissolved in sterile PBS. For intranasal (in) administration a volume of 50μL was employed. Sera samples, bronchoalveolar lavages (BAL), spleens and lungs were collected 15 days after the last dose.

### 2.3. Evaluation of humoral immune response by ELISA

The systemic and mucosal antibody response was evaluated by ELISA. Anti-IgG, subclasses and, -IgA ELISAs were conducted as previously described [**9–11**]. Briefly, 3 µg/mL of recombinant protein was used to coat 96 well high binding plates (Costar, USA). Plates were subsequently blocked with 2% skim milk solution. Samples were added in duplicates, starting from 1:100 dilution in the case of sera; whereas BAL were directly assayed. Specific horseradish peroxidase conjugates (Sigma, USA) were used. As substrate, OPD (Sigma, USA)/hydrogen peroxide solution was employed. Plates were incubated during 10 min in the dark and the reaction was stopped with 2 N Sulphuric acid. The optical density (OD) was read at 492nm in a multiplate reader (FilterMax F3, Molecular Devices, USA). The positivity cut-off was established as 2 times the average of OD obtained for a pre-immune sera pool. Data were represented as OD at 492 nm.

### 2.4. IFN-γ ELISPOT

IFN-γ ELISPOT assay was carried out using a specific antibody pair developed for the detection of this cytokine in mice samples (Mabtech, Sweden). Spleen and lung cells were isolated in RPMI culture medium (Gibco, US). Samples were processed individualized, with the exception of the placebo group for which a pool of three randomly selected mice was evaluated. Duplicates cultures (5x10^5^ splenocytes per well) were incubated for 48 h at 37°C, and 5% CO_2_, in a 96 well round bottom plate (Costar, USA) with a final concentration of 10 µg/mL of each stimulating agent: Ag85B peptides 10-27aa (1) (AWGRRLMIGTAAAVVLPG) and 91-108aa (2) (WDINTPAFEWYYQSGLSI), Ag85B protein, Concanavalin A (ConA) (Sigma, USA), or medium. The content of the plate was then transferred to an ELISPOT pre-coated plate, and incubated for 16-20 h at 37°C and 5% CO_2_. The successive steps were carried out following the manufacturers recommendations. Finally, the spots were counted using a stereoscopic microscope (AmScope SM-1TSZ, USA) coupled to a digital camera.

### 2.5. Transmission Electron Microscopy

For microscopy analysis, a specialized service was contracted with ETest company (Changsha, China). Briefly, samples of the recombinant protein Ag85B, in PBS, or mixed with ODN-39M or LAL_32-51_, were placed on a freshly glow-discharged, 400-mesh copper grid coated with formvar and carbon. After 2 min of sample absorption, the grids were washed with water and uranyl acetate stain was applied. Following 4 min of staining, grids were wick-dried using Whatman no. 1 filter paper and allowed to air-dry for 20 min.

A Transmission Electron Microscope HT 7800 (Hitachi, Tokyo, Japan) was employed for sample visualization. An acceleration voltage of 120 Kv and three magnifications, 25,000×, 50,000× and 100,000×, were used. Eight random fields per sample were photographed and analysed. The average particle size was estimated by digital measurement of the particle diameter using Image J software (version 1.4.3, Bethesda, MD, USA).

### 2.6. Study of NALT antigen-presenting cells after in vivo administration of synthetic adjuvants

Three Balb/c mice (6 to 8 weeks old) per treatment group received a dose of 20µg of ODN- 39M, or 2mM LAL_32-51_, dissolved in sterile PBS, or PBS alone, by intranasal route in a final volume of 20µL. For the administration, not anesthetized mice received the total volume very slowly, drop by drop, using a micro-pipette. Sixteen hours later the animals were sacrificed by cervical dislocation and the hard palate NALT-associated cells isolated as described by Cisney, E.D. et al, 2012 [**24**]. Briefly, the bell-shaped hard palates were excised using a No. 11 scalpel blade and gently pulled out with fine forceps. Then the palates were placed into a Petri dish containing supplemented RPMI culture media, washed twice and minced. Following, a pool containing the hard palates from 3 mice was incubated with 0.3mg/mL Collagenase D (Roche, Germany) during 1h in agitation. The obtaining solution was rinsed twice with culture media and the cells were counted. 5x10^5^ cells per condition were labeled for FACs analysis.

### 2.7. Flow cytometry

NALT cells were incubated 30 min at RT with Fc block (Innovex Biosciences, USA). Lymphocyte populations were detected after 20 min at 4℃ labeling with specific antibodies (proteintech, USA) for cell phenotyping and activation markers detection. PE-conjugated anti-mouse B220 antibody (clone RA3-6B2), PE anti-mouse CD11c (clone N418), CoraLite488-conjugated anti-mouse CD11b (clone M1/70), APC-conjugated anti mouse CD69 (clone H1.2F3), CoraLitePlus750-conjugated anti mouse CD103 (clone 2E7), FITC-conjugated anti mouse MHCII (clone M5/114.15.2), APC-conjugated anti mouse CD80 (clone 16-10A1), and CoraLite594 conjugated anti mouse CD86 (clone 230476B8) were employed. In addition, during the NALT isolation procedure establishment, the Alexa Fluor488-conjugated anti-mouse CD3 (clone17A2) (Invitrogen, USA) antibody was used to identify the lymphocyte population. All antibodies were diluted in FACS buffer (2% fetal bovine serum in PBS) at the manufacturer’s recommended concentration per number of cells. The samples were analyzed immediately after labeling, data were acquired with a FACs Melody (Beckton Dickinson, USA) and analyzed with FlowJo software (Tree Star, Ashland, OR).

### 2.8. Statistical analysis

Graph Pad Prism version 5.00 software (Graph-Pad Software, San Diego, CA, USA) was employed for the statistical analysis. One-way Anova test was used as parametric tests for multiple group comparisons, followed by a Tukey’s post-test. In the particular cases that required it, non- parametric multiple comparisons using Kruskal Wallis test and Dunns post-tests were employed. The statistical criteria followed the standard considerations for P values: ns, p>0.05; *, p<0.05; **, p<0.01; ***, p<0.001.

## 3. Results

### 3.1 Immunogenicity of Ag85B by intranasal route exploring two different synthetic adjuvants

In a first experiment we found that the Ag85B preparation when is administered by intranasal route in PBS is able to induce a specific IgG antibody response in sera of Balb/c immunized mice, even at the low dose employed of 10μg (Figure 1a). The use of the synthetic adyuvants evaluated, LALF_32-51_ peptide and ODN-39M, did not induce an increment in the antibody response measured in sera (IgG) neither in BAL(IgA) (Figure 1a and b, respectively). In the last case a slight increase in the specific IgA response in BAL was detected for the group receiving Ag85B+ODN-39M. Consistently, this group showed a significantly higher IFN-γ secretion response in spleens, compared with the control receiving the Ag85B alone (Figure 1c). The specific IFN-γ secretion detected at spleen, after the intranasal immunization, was positive only in response to the Ag85B protein stimulation. No positive response was observed for the Ag85B derived individual T CD4+ peptides evaluated (data not shown).

**Figure 1.**
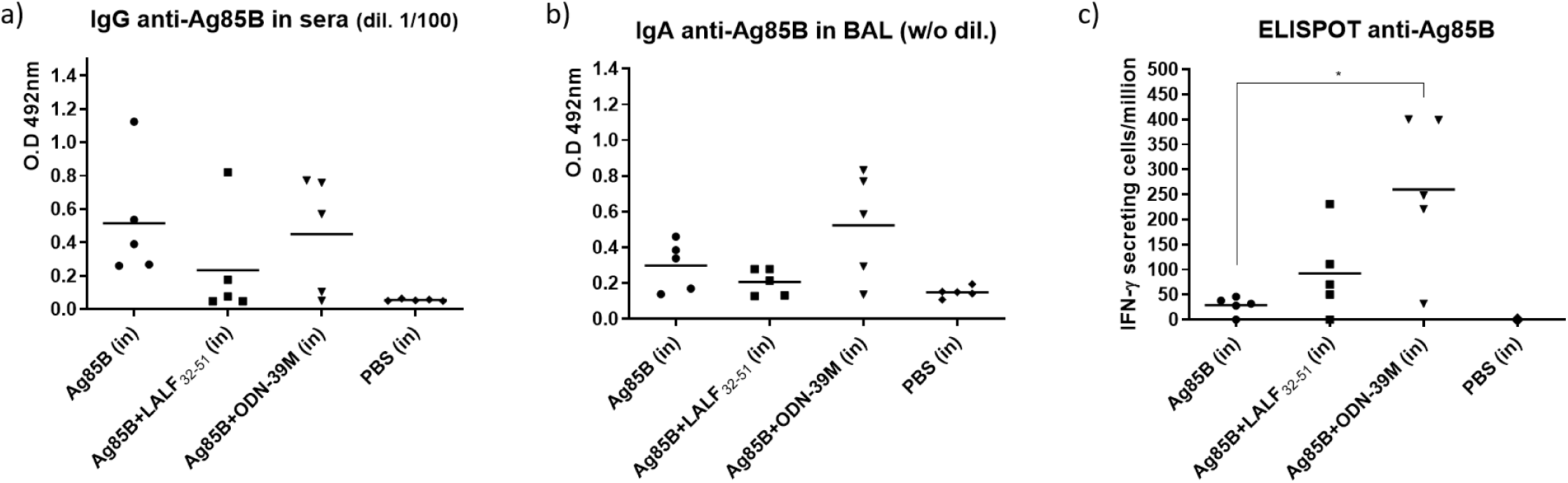
Immune response induced in Balb/c mice after intranasal administration of Ag85B (10μg) plus either LALF_32-51_ or ODN-39M as synthetic adjuvants. Mice received three doses on days 0, 15 and 30; and the samples were collected fifteen days after the last dose. The specific antibody response was evaluated by ELISA a) IgG in sera (diluted 1:100), and b) IgA in BAL (samples without dilution). c) IFN-γ secretion measured by ELISPOT in spleen. * means statistical significance p<0.05

A second study was done to further evaluated the immune response induced in lungs after the intranasal administration of the Ag85B preparations (Figure 2). In this second experiment a trend to induce a higher IgG response in sera was observed for the group receiving the Ag85B adjuvated with ODN-39M compared with the Ag85B in PBS (Figure 2a). To achieve a deeper characterization of the Ag85-specific antibody response generated after intranasal immunization with these preparations, the IgG1 and IgG2a subclasses were evaluated in the sera. Figure 2b shows that the group receiving the ODN-39M as adjuvant induced a remarkable high IgG2a response against the Ag85B, while for the group without adjuvant only IgG1 was induced. On the other hand, the IgA response measured in BAL only showed high levels for the group immunized with Ag85B+ODN-39M (Figure 2c). In line with this result, when we measured the IFN-γ secretion response elicited by lung resident lymphoid cells the same group of mice developed a positive response in the 3 out 3 animals tested, two of them with high levels (Figure 2e). Consistently, the two animals that developed the higher IFN-γ response in lungs coincided with the two that show also the higher response at spleen level (Figure 2d).

**Figure 2.**
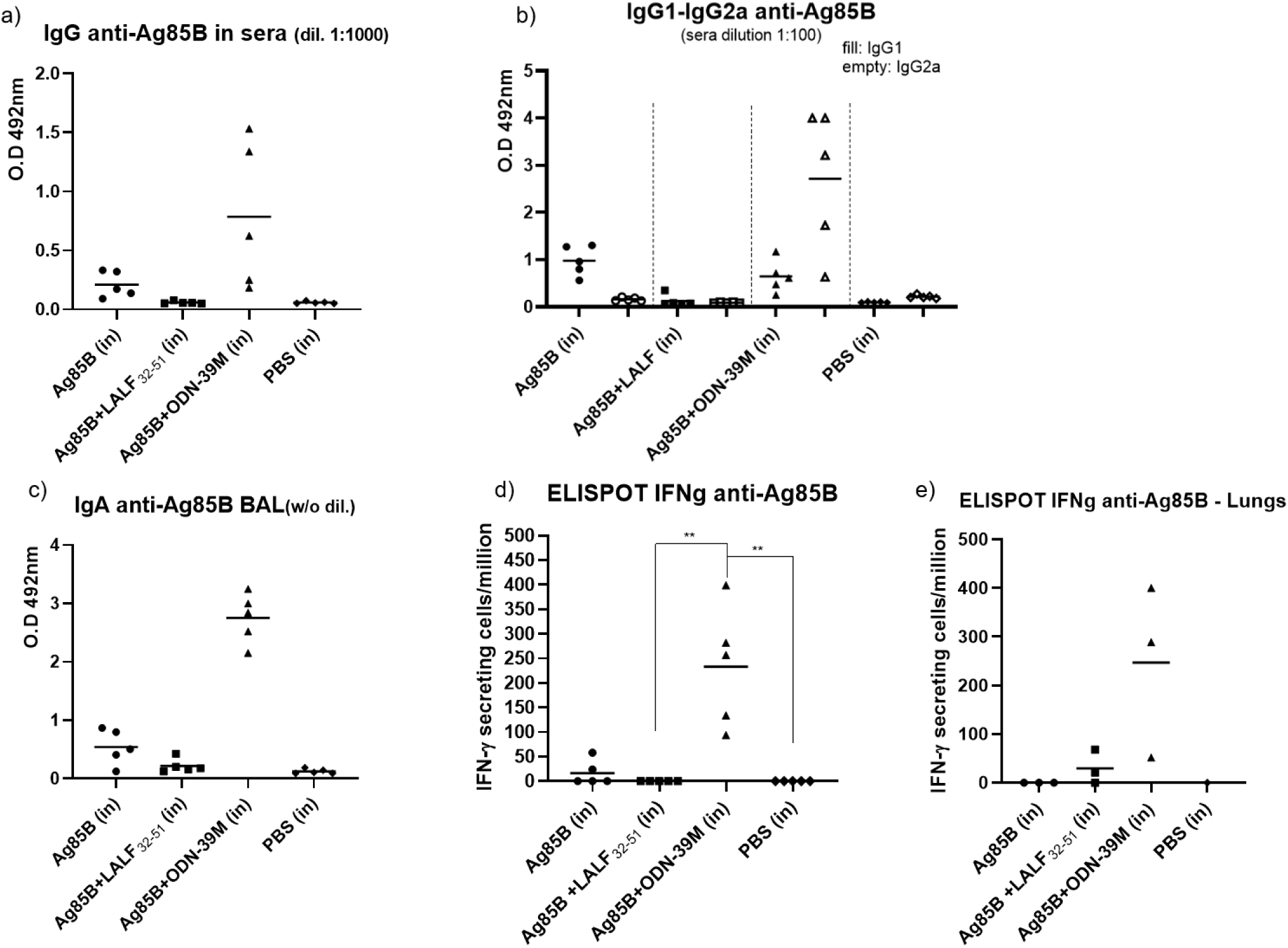
Immune response induced in Balb/c mice after intranasal administration of Ag85B (10μg) mixed with either LALF_32-51_ or ODN-39M as synthetic adjuvants. Mice received three doses on days 0, 15 and 30; and the samples were collected fifteen days after the last dose. The specific antibody response was evaluated by ELISA: a) IgG in sera (diluted 1:1000), b) IgG1 and IgG2a in sera (diluted 1:100), and c) IgA in BAL (samples without dilution). The cell-mediated immune response was evaluated by IFN-γ ELISPOT: d) in spleen cells, and e) in lung resident lymphoid cells. ** or * means statistical significance at p<0.01 or p<0.05, respectively.

In general, the immune response induced in this second animal study for the groups intranasally immunized with Ag85B in PBS, and adjuvated with ODN-39M, reproduce the data of the first study. However, for the group receiving the LALF_32-51_ peptide as adjuvant the results observed in the second experiment were more marginal in terms of specific IFN- γ secretion at spleen (Figure 2d), although a low but detectable response was found at lungs (Figure 2e).

### 3.2 Evaluation of Ag85B based vaccine preparations by Transmission electron microscopy

With the aim to obtain some clues regarding the potential mechanism of action of the vaccine preparations administered by intranasal route, we explore the structure of the different mix by TEM analysis. Previous data indicate that the mix of ODN-39M with nucleocapsid proteins from different virus promotes the formation of nanoparticles (NPs) and bigger protein aggregates [**9, 10, 17, 18**]. However, the present work constitutes the first evaluation of this specific CpG ODN combined with a bacterial derived recombinant protein. The results obtained after visualization of the Ag85B alone (dissolved in PBS) and in individual mix with ODN-39M or LALF_32-51_ peptide are shown in Figure 3.

**Figure 3.**
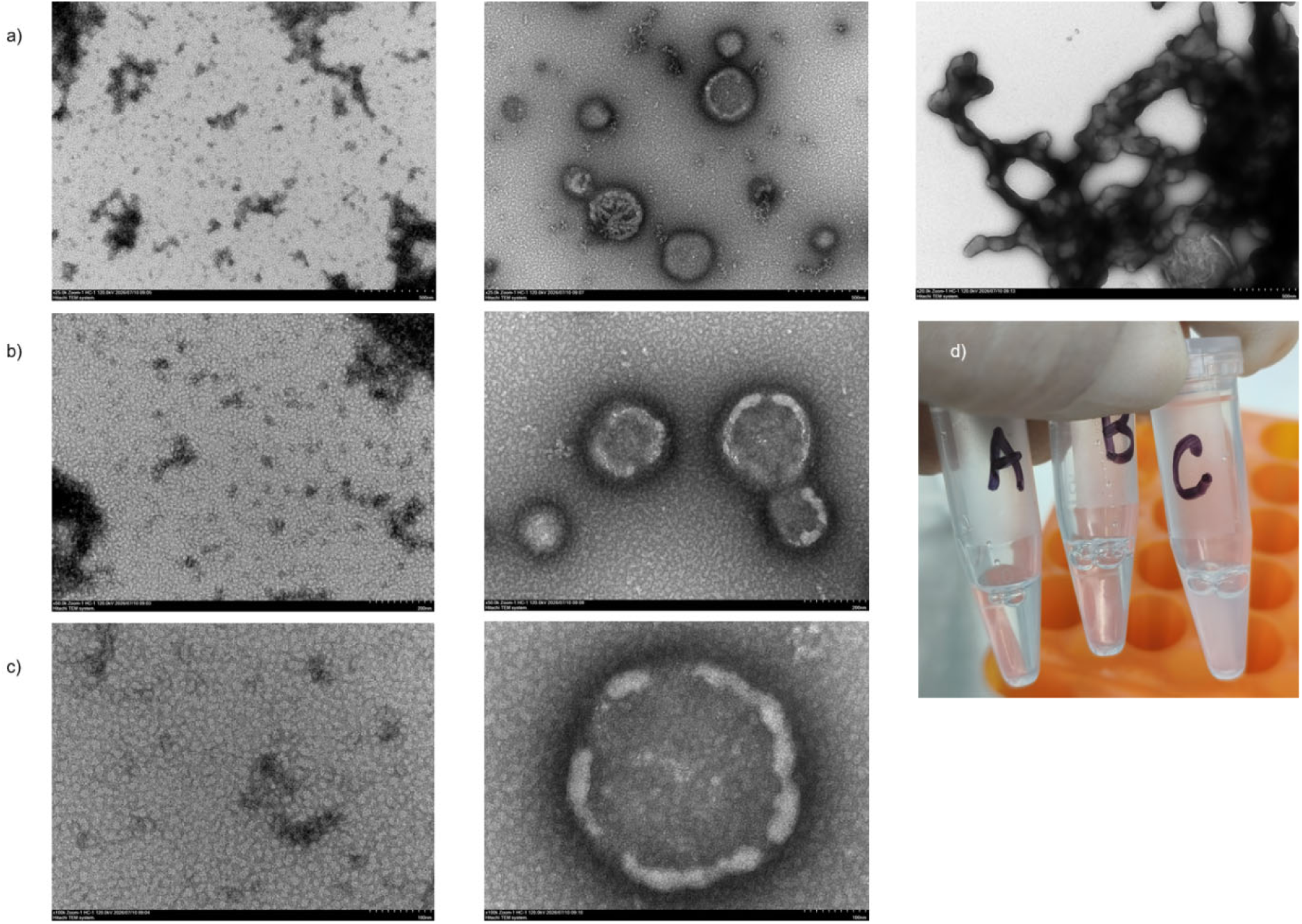
Transmission electron microscopy of the different Ag85B-based vaccine preparations. The samples were prepared using the same proportions employed in mice immunization experiments. For the visualization, a microscope Hitachi HT 7800 (voltage 120K) and three different magnifications x25.0k (row a), 50.0k (row b) and 100.0k (row c) were used. Ag85B dissolved in PBS (left panel), Ag85B + ODN-39M (middle panel) and Ag85B + LALF_32-51_ (right panel). For the last preparation only one magnification was applied. (d) Visual appearance of each preparation, A: Ag85B in PBS, B: Ag85B + ODN-39M, C: Ag85B + LALF_32-51_.

The formation of particles and aggregates of particles in the nanometre scale were observed for the Ag85B protein sample (Figure 3, left panel). Interestingly, when the recombinant Ag85B protein is mixed with ODN-39M, the formation of bigger spherical structures is observed (Figure 3, central panel), ranging around hundred of nanometres. These structures show a discontinuous bilayer surrounding delineating the spherical surface, and some of them present invaginations. On the other hand, for the sample of Ag85B + LALF_32-51_ peptide only one picture was available showing a micrometre size atypical structure (Figure 3, right panel).

### 3.3 Activation of NALT antigen-presenting cells after ODN-39M or LALF_32-51_ *in vivo* intranasal administration

With the aim to study the local mechanism of action of the synthetic compounds under evaluation as nasal vaccine adjuvants, we implement the isolation of nasal associated lymphoid tissue immune cells following the method published by Cisney, E.D. et al, 2012 [**24**]. Several experiments were carried out to establish the methodology in our lab. To identify the target cell population, we used FACs with specific fluorophore-labeled antibodies for phenotyping the cells. Figure 4 shows a representative assay isolating in parallel lymphoid cells from NALT and spleens from Balb/c naive mice. To our knowledge very scarce information has been published focusing on immune cells isolated from mice NALT tissue. In line with the previous documented experience [**24**] a very low yield of immune cells is obtained from NALT per mouse, indicating that pooled samples, from at least three mice per treatment group, should be used for subsequent experiments.

**Figure 4.**
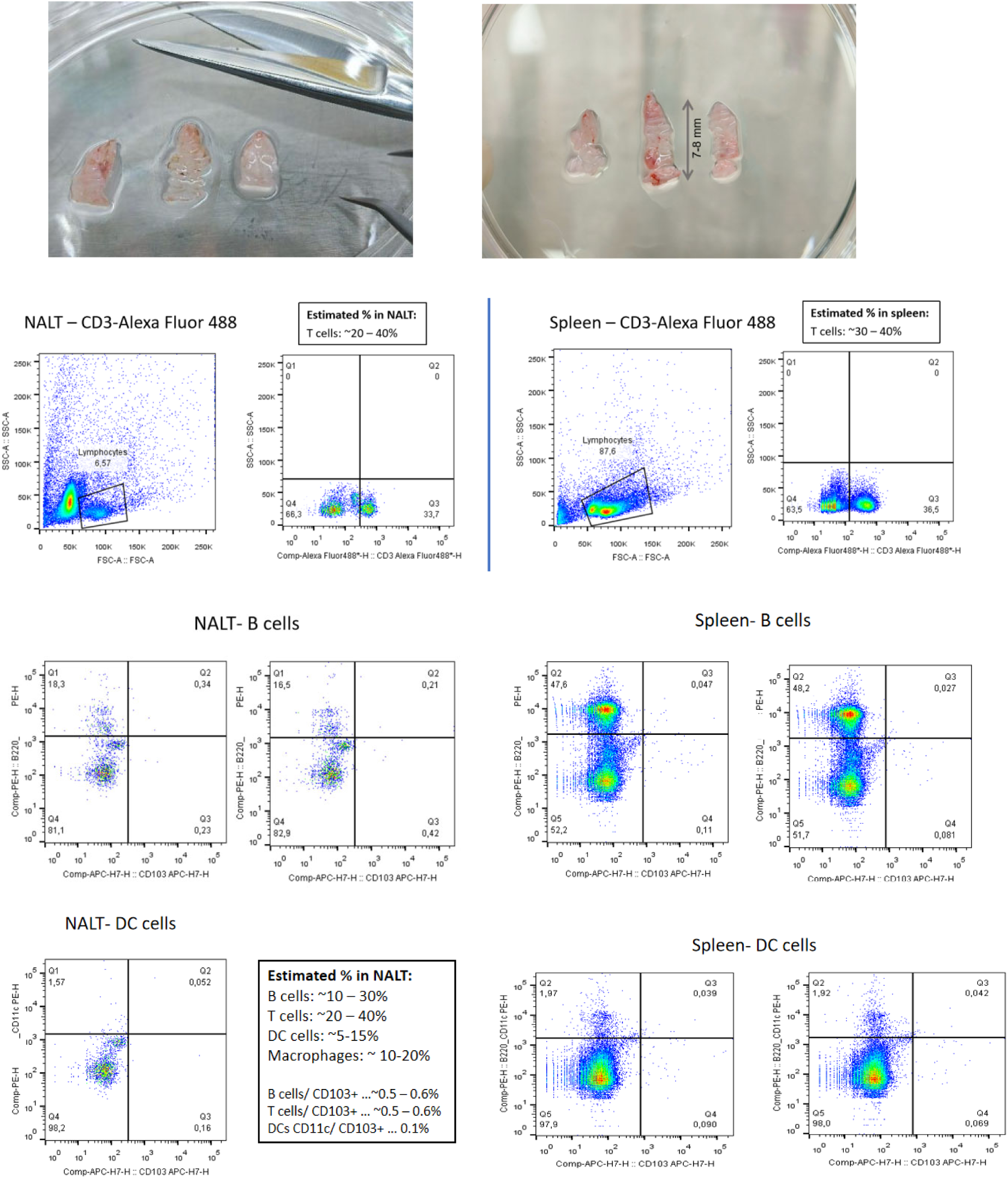
NALT derived immune cells isolation. Representative hard palate samples collection as the departure biological material to isolate the immune cells associated to nasal passage (superior panel). The lower panel shows the FACs plots representing the identification of T, B and dendritic cell (DC) cell populations in NALT samples (left), and spleen samples (right). For the identification of each specific cell population the following antibodies were employed: Alexa Fluor488-conjugated anti-mouse CD3 (clone17A2), PE-conjugated anti-mouse B220 antibody (clone RA3-6B2), PE anti-mouse CD11c (clone N418). Small squares with the previous reported percentage of each cell population at NALT location are also included in the figure.

Considering the promising results obtained using the ODN-39M as adjuvant by intranasal route we decide to preliminary explore which kind of antigen presenting cells are activated in the NALT after ODN-39M administration to mice. Sixteen hours after intranasal administration of ODN-39M we observed a marked increase in the total number of cells present at NALT tissue compared with PBS treated animals (Figure 5).

**Figure 5.**
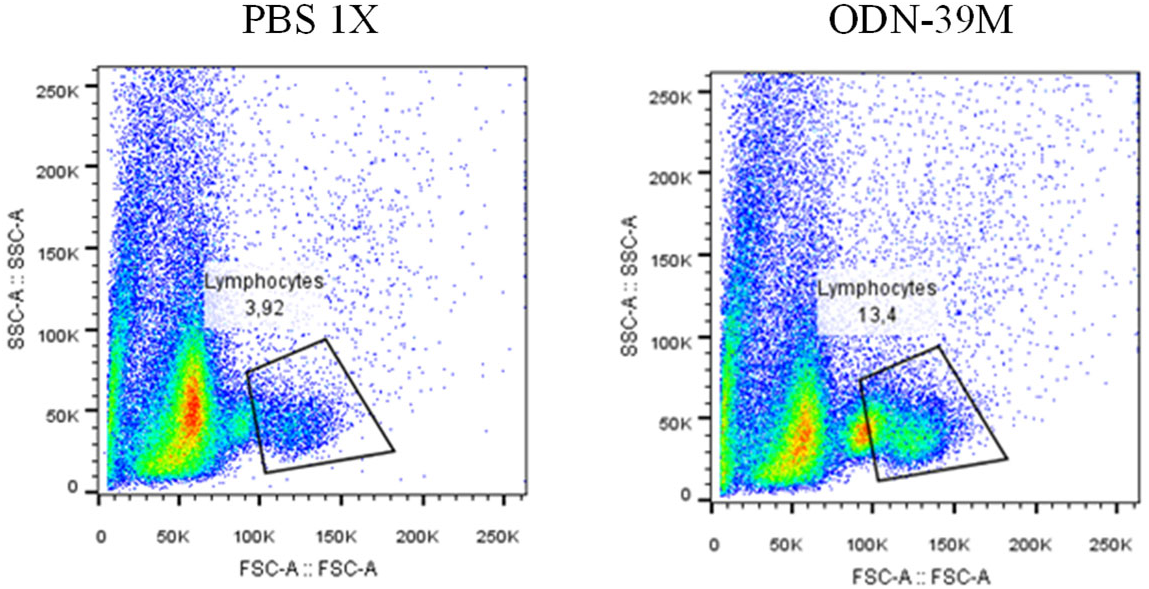
Isolation of hard palate associated NALT. Three mice per groups were intranasally immunized with 20ug ODN-39M or PBS1X. Sixteen hours later the animals were sacrificed and the NALT tissue immune cells isolated for FACs analysis. The gating represents the lymphoid population. Mice treated with PBS1X (left) and ODN-39M (right).

The B cells and dendritic cells (DCs) found in NALT, show a notable increase in their expression of CD69 early activation marker after intranasal administration of ODN-39M (Figure 6). On the other hand, a preliminary experiment was also carried out to evaluate the early effect of LALF_32-51_ peptide in mice NALT, under similar assay conditions. In this case, no remarkable increase in the immune cell’s recruitment was detected for the peptide treated sample. However, the results show a moderate increase of CD69 marker in B cells and in myelod derived immune cells (CD11b+ population), after LALF_32-51_ peptide stimulation (Figure 7). For both stimulus other activation markers like MHCII, CD80 and CD86 were also evaluated but no clear increase in their expression was detected. In addition, the expression of CD103 surface marker, associated with mucosal tissue retention of immune cells, was studied but the levels detected were marginal in all our experiments.

**Figure 6.**
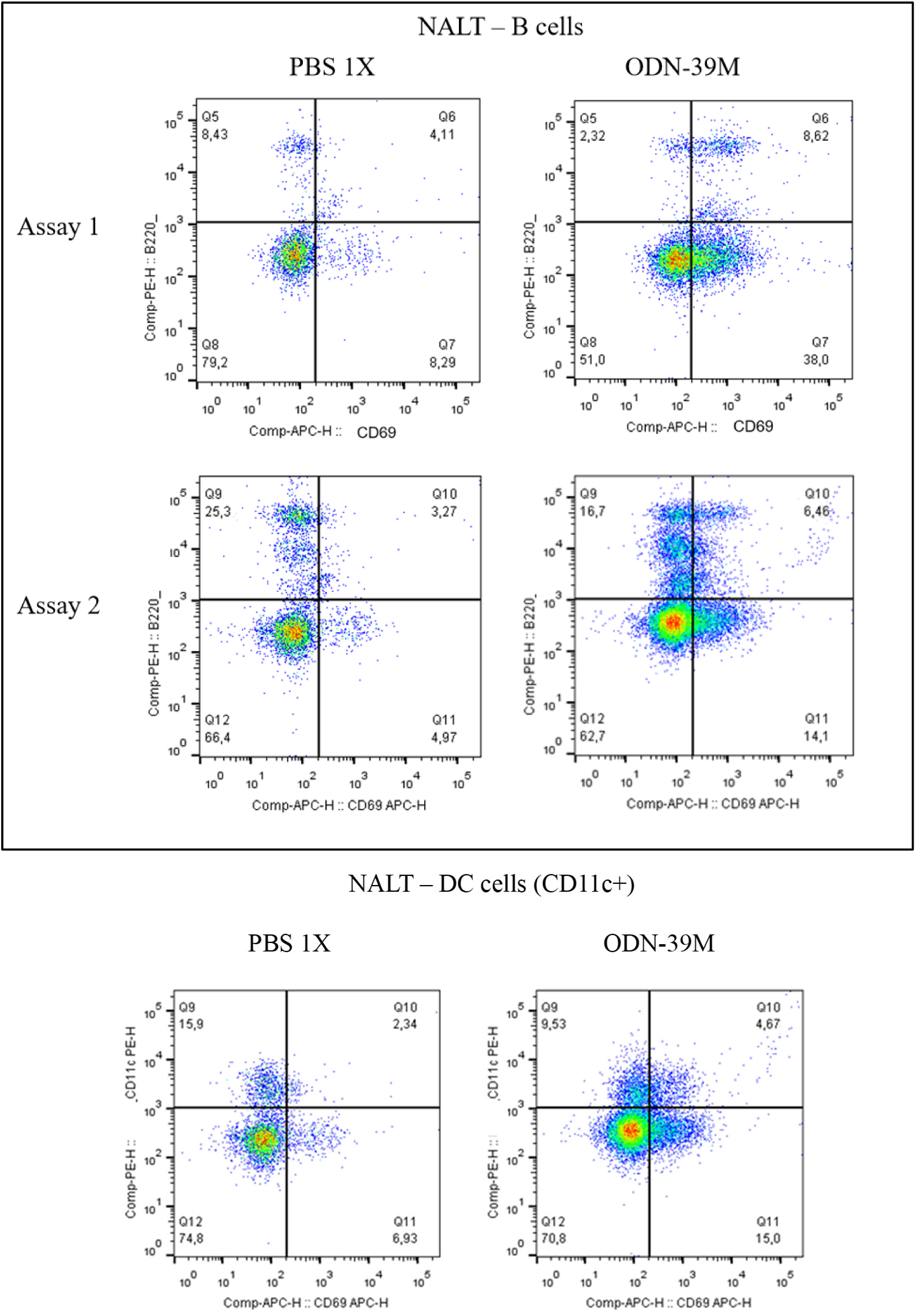
Expression of CD69 early activation marker in NALT immune cells. Three mice per groups were intranasally immunized with 20ug ODN-39M or PBS1X. Sixteen hours later the animals were sacrificed and the NALT immune cells isolated and labeled for FACs analysis. Upper panel represents B cells (labeled with anti-B220-PE) analyzed in two independent assays. Lower panel represents CD11c+ cells (labeled with PE anti-mouse CD11c (clone N418) coming from one representative assay. Dot plot representations and population estimations at each analysis quadrant using FlowJo. 20,000 cell counts were acquired for the analysis.

**Figure 7.**
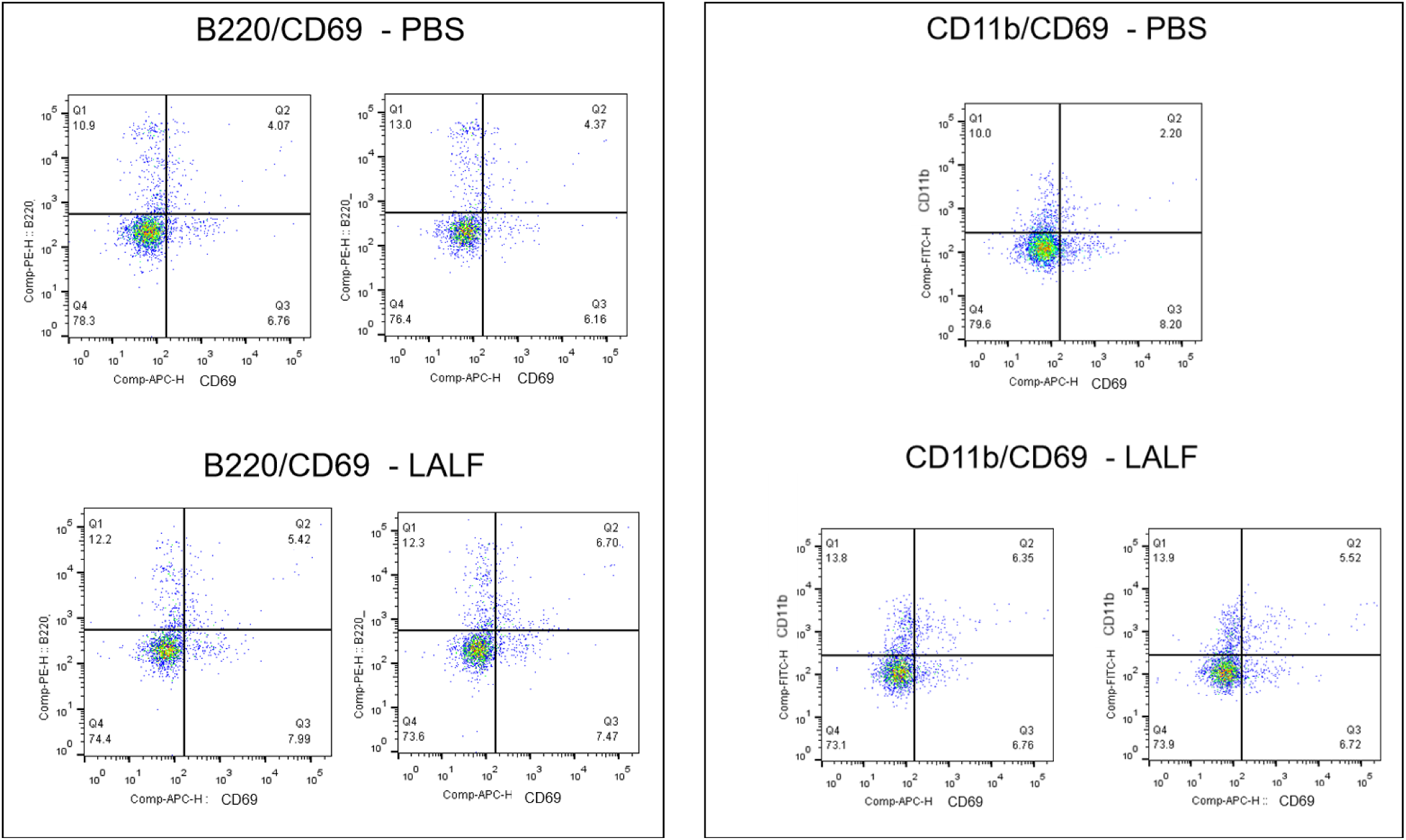
Expression of CD69 early activation marker in NALT immune cells. Three mice per groups were intranasally immunized with 2mM LALF_32-51_ or PBS1X. Sixteen hours later the animals were sacrificed and the NALT immune cells isolated and labeled for FACs analysis. Left panel represents B cells (labeled with anti-B220-PE). Right panel represents CD11b+ cells (labeled with CoraLite488-conjugated anti-mouse CD11b (clone M1/70). Dot plot representations and population estimations at each analysis quadrant using FlowJo. 10,000 cell counts were acquired for the analysis.

## 4. Discussion

In this work we evaluated the adjuvant properties of two synthetic compounds over a model TB antigen following intranasal administration of their single mix ie., antigen+adjuvant, in Balc/c mice. The Ag85B was selected as target antigen considering that is one of the most dominant protein antigens secreted from all mycobacterial species and has been shown to induce substantial Th cell proliferation and vigorous Th1 cytokine production in humans and mice. Actually, Ag85B constitutes the base of several vaccine candidates under evaluation nowadays [**6, 7**]. The Ag85B possesses mycolyl-transferase and fibronectin binding activity [**25**]. This last, facilitates the attachment of Mtb to alveolar macrophages. It is known that fibronectin glycoprotein, a main component of the extracellular matrix (ECM) proteins is present in mucosal surfaces [**26**]. Specifically, in murine models, this protein is constitutively expressed in the basement membrane and subepithelial layers of the respiratory mucosa, including the nasal passages. Fibronectin can be considered moderately abundant in the murine nasal mucosa, especially during tissue remodeling or immune responses. In addition, it has been detected in nasal respiratory and olfactory epithelium regions in mice [**27**]. Furthermore, some studies have shown that some ECM proteins, including fibronectin, can acquire properties of damage-associated molecular patterns (DAMPs) after modifications induced, at mucosal sites, by different agents, for example by bacteria or host proteases [**28**]. These findings could explain our results showing that recombinant Ag85B, without any additional adjuvant, is able to induce a detectable immune response after intranasal administration in mice. Altogether, the data suggest that Ag85B per se could activate in some extent the APCs at nasal mucosa, favoring the development of an immune response.

On the other hand, Ag85B contains multiple epitopes that could elicit humoral as well as Th1 or Th17 cellular immune responses, which provides a long-term protective response against M. tuberculosis infection [**29**]. The results obtained in the current work are in line with the features previously described for the Ag85B protein. Our data indicate that the Ag85B protein preparation employed, without adjuvants, was able to induce an immune response after intranasal administration, even at the relative low dose used of 10μg per mice. In the two independent experiments carried out, the group of mice receiving the Ag85B in PBS showed a low, but positive, IgG antibody response in sera, IgA in BAL and IFN-γ secretion at spleen level. It is recognized in the literature that soluble proteins do not result immunogenic after intranasal administration, requiring the use of more complex formulations or potent adjuvants [**30**]. Considering this, in our opinion the Ag85B protein has intrinsic features that favor its immunogenicity by intranasal route. This behaviour could be related with the Ag85B protein binding capacity to fibronectin, as mentioned above, but also with the formation of particles and aggregates in the recombinant Ag85B preparation employed as immunogen. The particulate nature is not intrinsic to this protein’s native biological state, but can arises during is recombinant expression. Actually, results coming from Ag85B SDS-PAGE evaluation under native and reduction conditions show the purified recombinant Ag85B used in our studies can form aggregates of high molecular weight, consistent with dimer size (data not shown).

Furthermore, the particulate/aggregate structure of the used preparation of Ag85B in PBS was visualized by TEM. Surprisingly, for the preparation of Ag85B + ODN-39M the formation of bigger aggregates structures resembling Mtb bacteria were observed. The spherical structures have a diameter size in the range of hundred nanometres and for some of them a discontinuous bilayer with invaginations surround the sphere surface. The particular structure observed for the ODN-39M adjuvated preparation can explain the high immunogenicity found for this combination after intranasal administration in mice. On the other hand, the preparation containing Ag85B + LALF_32-51_ peptide shows a complex and aberrant structure with a micrometre size. The advantages of particulate / aggregates antigenic preparations, in the nanometre range, on vaccines immunogenicity are well recognized, being a more relevant feature for nasal administered vaccines. This kind of structures promote the up-take by antigen presenting cells, favoring the induction of a stronger immune response compared with soluble proteins. In general, the current TEM findings correlate with the immune response observed for each preparation, and are in line with previous data supporting the important role of particle size in mucosal administration. Studies using polystyrene NPs in human respiratory mucus has shown that particles with a size of 100 nm and 200 nm can quickly penetrate into respiratory mucus, while particles with a size >500 nm remain blocked and can’t pass through the mucus barrier. Other data indicate that the effective diffusion rate of NPs in mucus decreases with the increase of particle size (20–500 nm). Larger drugs (usually in the micron range) cannot diffuse in the mucus layer due to their steric hindrance [**31**]. For the preparation of Ag85B + ODN-39M, the effect of a proper nanoparticle size in addition to the potential favored interactions between this specific protein with fibronectin in ECM, and the potent immunostimulatory capacity of the CpG ODN; could jointly explain the better immunogenicity results.

The evaluation of the two potential mucosal adjuvants in study using the Ag85B as TB model antigen by intranasal route, clearly confirm the ability of the CpG ODN, ODN-39M, as enhancer of the immune response with a bias to develop a Th1 pattern. The obtained results are in line with previous data evaluating the ODN-39M as nasal vaccine adjuvant using different recombinant antigens from SARS-CoV-2 [**9–11**]. However, on the other hand, the LALF_32-51_ peptide which previously has shown an immunoenhancing effect over the HBsAg, and an HPV-derived, recombinant protein after intranasal administration in mice [**32**], did not induce a similar effect when was evaluated with the Ag85B in the present work.

LALF_32-51_ is a primary amphipathic peptide, previously described as an immunomodulatory peptide with cell-penetrating capacity [**33–35**]. Based on these properties the peptide has been used in the design of cancer therapy and an HPV vaccine candidate for parenteral administration [**34–36**]. The use of CPPs in vaccine formulations promote the antigen direct translocation into the cytosol of APCs, thereby accessing the subcellular compartments where classical MHC class I peptide loading occurs. This approach would enable not only cross-presentation competent cells to present vaccine antigens on MHC-I, increasing the priming of CD8+ T cells. On the other hand, the charge distribution of the LALF_32-51_ molecule (with highly cationic regions) could favor its interaction with the mucin protein, the main component from the mucus, which is negatively charged [**37**]; increasing the residency time of the vaccine preparation at the nasal mucosa. This behavior has been reported for other compounds, like chitosan nanoparticles, administered by nasal route [**38**]. However, a light turbid appearance was observed in the Ag85B+LALF_32-51_ preparation, potentially related with electrostatic interactions, that could be affecting the effect of LALF_32-51_ over this specific antigen. The formation of micrometer-size aggregated structures, visualized by TEM, can be an effect of the electrostatic interactions, and affects the development of a potent immune response for this preparation after nasal administration. In this line, the negative results obtained for the LALF_32-51_ peptide in the current work seem to be formulation related, therefore its immunoenhancing capacity by nasal route over other antigens can not be discarded and should be explored case by case.

On the other hand, a recent publication describes the evaluation of another synthetic adjuvant, c-di-AMP, along with Ag85B by intranasal route in a Mtb persistent infection mouse model [**39**]. The results showed by this group suggest the potential of c-di-AMP in the generation of immune response and protective effect at lung level in a therapeutic scenario. This work emphasized the relevance of looking for new mucosal adjuvant options for TB new vaccine developments. However, the authors employed a dose of 30μg of Ag85B per mice, which represents three time more than the dose used in our experiments. Usually, the studies published using the Ag85B, and other related TB protein antigens, in mice models employ parenteral administration with doses between 20 to 50μg per mice, and in many cases the animals are pre-sensitized with BCG immunization [**40, 41**]. In our previous research using protein subunit immunogens from Coronavirus and other pathogens (mainly viruses), by nasal route, we consider the dose of 10μg per mice a moderate to low level dose, suitable for mice studies [**9–11, 32**]. The dose of Ag85B selected for the current experiments was also constrained by the availability and price of the purified recombinant protein. Although we found appealing results using the dose of 10μg per mice by intranasal route, in our opinion higher doses could improve the observed immune response. In addition, further studies employing more specific TB animal models, or BCG-primed mice, are recommended for the Ag85B+ODN-39M nasal vaccine preparation.

In parallel, understanding the cellular and molecular landscape of the NALT has significant translational implications for nasal vaccine development. However, despite the growing interest in using the intranasal route for mucosal vaccine delivery, modest attention has focused on the local site(s) and mechanism(s) that might be responsible for the induction of such responses. In rodents, NALT comprises a pair of lymphoid aggregates located on the either side of the nasal passages, above the hard palate. The structural and functional evaluation of NALT in mice, and its homologous structure in humans (the Waldeyer’s ring and the adenoid, tubal, palatine, and lingual tonsils), has been challenging, largely because of the complexity in accessing its anatomic location and also by the low yield of immune cells that can be recovered [**42**].

In order to elucidate the mechanisms of action of the evaluated adjuvants after intranasal administration in mice, the activation of NALT immune cells was preliminary studied using flow cytometry. In our activation experiments, we measured the expression of CD69, CD80, CD86 and MHCII in NALT more prominent antigen presenting cells. In both DCs and B cells, CD69 functions as a highly sensitive, transient indicator of early immune activation. CD69 expression can be up-regulated very fast in response to stimulus, between 3-4 hours. Its expression remains highly detectable at around 15 hours post-stimulation, then a general decrease occurs from 24 hours on [**43**]. Some differences on the time of CD69 expression up-regulation onset has been observed, depending on the nature of the stimulus, for example, CpG has been shown to induce faster B cell activation based on CD69 expression compared to LPS (a TLR4 ligand) [**44**]. On the other hand, CD80, CD86, and MHCII are core maturation and antigen-presentation molecules, critical for T cell activation, with distinct temporal profiles. CD86 is the initial co-stimulatory ligand, with an earlier expression pattern compared to CD80. In many murine models, CD86 is detectable within hours of stimulation, often peaking between 12 to 24h. While CD86 peaks early, CD80 levels can continue to rise and be boosted late, some studies show increased CD80 surface levels even after 60h post-stimulation [**45**]. In contrast, to the previous mentioned markers, which are induced from very low basal levels, MHCII is constitutively expressed at moderate levels on mature DCs and B cells, activation simply up-regulates the existing expression. MHCII up-regulation can happens rapidly, some studies show a plateau around 8h post-stimulation, remaining high for an extended period from 12h and beyond [**46**]. Taking into account the above-mentioned reports, documenting the kinetic of expression of these relevant APCs surface activation markers, in our experiments we chosen a stimulation time of 16h after *in vivo* intranasal administration.

From previous works it is known that CpG ODN are able to bind and activate toll-like receptor 9 (TLR9) present at antigen presenting cells [**47**]. A similar mechanism of action is expected for the ODN-39M, which is a complete phosphodiester backbone ODN designed with motifs able to active mice and human immune cells. Furthermore, Pohar J et al [**48**] described an increase of TLR9 activation by phosphodiester backbone CpG motif ODNs. This kind of CpG ODN also has the advantage of a potential safer profile compared with the more extended phosphorothioate backbone variants, which has been identified as a factor contributing to the adverse reactions detected in therapeutics interventions [**49**].

TLR9 is expressed mainly by B cells and dendritic cells (DCs) [**50**], both playing key roles in innate/mucosal immunity. NALT lymphoid compartment contains the typical cell populations of an immune-inductive site, specifically enriched in plasmacytoid dendritic cells, which are activated by TLR9 agonists, as the CpGs ODNs [**51**]. In line with this data, a considerable higher number of lymphoid cells in ODN-39M-stimulated mice NALT samples were found compared with PBS receiving mice, this could be a direct effect of the stimulus in the recruiting of immune cells. Altogether, contributing to the activation of local immune system, favoring the development of an adaptive response to vaccine preparations. Furthermore, the CD11c+, and B220+ cells present at NALT of animals receiving ODN- 39M show a marked increase in the expression of the CD69 activation marker compared with PBS stimulated animals. CD69 expression is readily upregulated upon activation in most leukocytes, underlying its widespread use as early marker of activated lymphocytes [**43, 52**]. CD69 also plays a functional role in determine the patterns of cytokine release as well as the homing and migration of activated lymphocytes, promoting the retention of lymphocytes in tissues, including mucosal sites like the nasal epithelium. Correlating with the present results, a recent study found the up-regulation of CD69 activation marker in airway immune cells following intranasal vaccination [**53**]. Furthermore, in line with our results, Nacer A et al [**54**] found that NALT from mice immunized with flagellin (a TLR5 agonist) -containing preparations were characterized by and expansion of B and T lymphocytes and an *in vivo* influx of CD11+ DCs, suggesting key roles for these cells in nasal vaccine-induced immune responses. In addition, they observed that the majority of lymphocytes in the naive NALT were B220+ B cells. This work and previous studies show that NALT experiences an early T and B cell expansion in response to immunization, followed by draining lymphonodes later expansion.

On the other hand, considering previous studies on macrophages-derived cells documenting that LALF_32-51_ has immunomodulatory effect and significantly increases mRNA levels and cell surface expression of TLR2 and TLR4 [**33**] we decided to explore, in addition to B cells, other cell populations in the NALT from animals treated with LALF_32-51_. As TLR2 and TLR4 are most highly represented in cells of myeloid origin, which act as primary sentinels detecting invading pathogens, we selected the CD11b+ cell subset. CD11b surface marker is a broad recognized pan-myeloid marker found on almost all innate immune cells (neutrophils, macrophages, monocytes, DCs) [**55**]. Consistent with the expected, 16h after LALF_32-51_ intranasal administration, we found a moderate increase in CD69 activation marker for the two cell populations evaluated. However, for none of the two evaluated adjuvants we detected a significant increase in other activation markers (CD80, CD86, MCHII), which could be related with the time point selected for the study or with the fact that we evaluated the adjuvants alone, without including antigens in the intranasal administered preparations. More detailed evaluations are ongoing exploring different stimulation conditions. On the other hand, in line with our results, some authors have found that murine NALT contains a low frequency of CD103+ DCs [**56, 57**].

Interesting it has been reported that within the murine NALT, B cells are predominant and also a high density of CD11c+ cells were found uniformly distributed on NALT surface [**56**]. Additionally, few works describe that some nasal DCs has been found extending dendrites across the nasal epithelium, into the lumen of the nose, for antigen sampling [**56**]. On the other hand, migratory behavior is a recognized hallmark of DCs. The antigen- induced activation and following migration of NALT DCs to draining lymphonodes, mainly cervical, has been previously documented [**56**]. In parallel, a recent report describes the cell composition from human NALT samples [**58**]. The pharyngeal tonsils (also known as adenoids), the major NALT site in humans, were accessed using a nasopharyngeal biopsy technique. Consistently with this tissue in mice, in humans there is a predominance of germinal center B cells and Tfh (T follicular helper) cells, and 11 mononuclear phagocytes cell types, leading by conventional and plasmacytoid dendritic cells, among others. In correspondence, the inclusion of a DC-specific nasal adjuvant constitutes a feasible strategy to induce a potent initial local response to nasal vaccines, which contributes to the development of effective and long-lasting adaptive immunity.

Summarizing, the current experiments focused on the study of NALT are just a preliminary approach, further studies have to be done to expand the understanding of the local mechanism of action of ODN-39M, LALF_32-51_ peptide, and other synthetic adjuvants following intranasal immunization. In this sense, other surface activation markers and cell populations, as well as effector response mediators (ex/ secreted cytokines and chemokines) must be considered. In addition, the adjuvant principle should be evaluated with and without the antigen, and at different relevant time points. On the other hand, the results obtained for the vaccine preparation containing Ag85B + ODN-39M are promissory. The development of Th1-biased T-cell responses, both in lung and systemic compartments, is widely recognized as critical for protection against tuberculosis [**59**]. Currently there are several new TB vaccine candidates under clinical and preclinical evaluation, several including Ag85B as target, however the present candidate stands out by combining very attractive features (ex/ nasal administration, VLP protein-subunit platform, high immunogenicity and expected safety profile) deserving further development. The Ag85B + ODN-39M vaccine preparation constitutes an appealing option to boost the immunity initially primed by the traditional BCG vaccine in adolescents and adults.

## Conclusions

The results coming from the evaluation of the ODN-39M as nasal vaccine adjuvant employing a formulation containing the Ag85B recombinant protein as antigen corroborate previous data indicating the potential of this synthetic ODN as a safe nasal immune- enhancer with a bias to develop a potent Th1 immune response. In mice the evaluated vaccine preparation generates humoral and cellular immune response at systemic and lungs level, resulting appealing for further evaluation in TB mice models and challenge experiments. In addition, some preliminary data were generated evidencing an effect of the ODN-39M in the recruiting of lymphoid cells to NALT tissue and its preferential early activation of dendritic and B cells. Altogether, the results pinpoint the ODN-39M as a promising adjuvant for other nasal vaccines developments.

## Declarations

### Ethics approval and consent to participate

Clinical trial number: not applicable. The animal study protocols were approved by the Institutional Animal Care and Use Committee at Beijing Vital River Laboratory Animal Technology Co., Ltd. The standard of laboratory animal room complied with the national standard of the people’s Republic of China GB14925-2010. Protocols: P202506180001 (approved on June 2025) and 202510100001 (approved on September 2025).

### Availability of data and material

We agree to make the data accessible in case of any specific request.

### Consent for publication

All authors have consented to the publish the manuscript.

### Author Contributions

Conceptualization, Y.L. and Y.P.; Supervision, Y.P.; Investigation, R.C., Y.L., T.C.; Formal analysis, Y.L.; Funding acquisition, Y.L.; Writing—Original Draft, Y.L.; Writing—Review & Editing, Y.L., Y.P.; Project administration, Y.L., W.L. and Y.P. All authors have read and agreed to the submitted version of the manuscript.

## Acknowledgement

Not applicable.

